# BASILIScan: a tool for high-throughput analysis of intrinsic disorder patterns in homologous proteins

**DOI:** 10.1101/378018

**Authors:** Michał S Barski

## Abstract

**Background:** Intrinsic structural disorder is a common property of many proteins, especially in eukaryotic and virus proteomes. The tendency of some proteins or protein regions to exist in a disordered state usually precludes their structural characterisation and renders them especially difficult for experimental handling after recombinant expression.

**Results:** A new intuitive, publicly-available computational resource, called BASILIScan, is presented here. It provides a BLAST-based search for close homologues of the protein of interest, integrated with a simultaneous prediction of intrinsic disorder together with a robust data viewer and interpreter. This allows for a quick, high-throughput screening, scoring and selection of closely-related yet highly structured homologues of the protein of interest. Comparative parallel analysis of the conservation of extended regions of disorder in multiple sequences is also offered. The use of BASILIScan and its capacity for yielding biologically applicable predictions is demonstrated. Using a high-throughput BASILIScan screen it is also shown that a large proportion of the human proteome displays homologous sequences of superior intrinsic structural order in many related species.

**Conclusion:** Through the swift identification of intrinsically stable homologues and poorly conserved disordered regions by the BASILIScan software, the chances of successful recombinant expression and compatibility with downstream applications such as crystallisation can be greatly increased.

## Introduction

The incidence of intrinsic disorder – defined as the lack of a fixed three-dimensional conformation of a protein - has become increasingly appreciated and examined in recent years. Many such proteins have been characterised, and their disorder, as well as order-disorder transitions upon ligand and partner binding demonstrated with a range of biophysical techniques (1). It is now widely acknowledged that protein disorder is a remarkably common phenomenon – especially in complex organisms and viruses. An estimated 30-40% of the human proteome is disordered to a significant degree (2). This has far-reaching consequences for the structural characterisation and experimental handling of many proteins from the human and other disorder-enriched proteomes.

Depending on the extent and location of flexibility, the intrinsically-disordered protein (IDP) can exhibit a number of different behaviours. Although the highly-charged nature of disordered regions may confer high solubility (3), presence of proteases in IDP preparations often causes severe proteolytic degradation in the case of fully-disordered IDPs or digestion of long connecting loops within and between otherwise structured domains (4, 5). Ultimately, biochemical and biophysical characterisation of such proteins is usually difficult because of challenging protein expression and purification and lack of sample homogeneity. Crystallisation of IDPs for X-ray crystallography is only feasible with the flexible regions removed, bound to a co-factor, or entropically-stabilised otherwise. Multidimensional nuclear magnetic resonance (NMR) spectroscopy remains the only high-resolution biophysical technique for studying some disordered protein systems, although the sample has to meet stringent compatibility standards (small and globular proteins, homogeneity, stability under low ionic strength for long periods of time) (6).

Since IDPs display characteristic patterns of amino acid content and distribution, the presence of disordered regions can be predicted from their primary sequence with high confidence. Many predictive algorithms have been devised and used successfully to predict the probability of residues in a given protein sequence to exist in an ordered or disordered state (7-11). Such predictions have been repeatedly confirmed experimentally with multidimensional NMR (12, 13). It is currently becoming common practice to take the intrinsic disorder predictions into account while designing a protein construct for recombinant expression. The N- or C-terminal disordered tails/domains can be truncated (for example: (14, 15)), while long loops connecting neighbouring domains or elements of secondary structure can be shortened to aid conformational stability (16). Unfortunately, the thin line between limiting disorder and affecting the function and correct folding of the protein can easily be crossed and the experimental trial-and-error process often takes long before an improved construct is found.

The software presented here, called BASILIScan, offers an alternative approach to streamline and simplify the construct design process. The core mechanism relies on a BLASTP search with the user’s amino acid sequence linked with simultaneous intrinsic disorder prediction of all hits. A specialised scoring system, called the FLEX score, is then employed to identify closest homologues exhibiting lowest disorder content. A hit possessing a FLEX score superior to the submitted protein is very likely to also show improved *in vitro* behaviour. Analytical features of BASILIScan, such as multiple sequence disorder overlay and alignment, allow for analysis of disorder conservation patterns between selected homologous hits in parallel. In addition to identifying a more suitable homologue, such analysis can guide rational construct design by exposing intrinsically-disordered regions of low or limited conservation which could be truncated from the expression construct. The adjustability of numerous search and analysis parameters provides compatibility with protein sequences of a wide range of intrinsic disorder, length and inter-species sequence variation.

## Design and Implementation

The main framework of the BASILIScan software is centred upon the connection of the modules outlined in Figure 1, written in Python 2.7. Where possible, Biopython (ver. 1.67) (17) libraries or their derivatives are used. In order to promote the multi-disciplinary use of BASILIScan, a graphical user interface (GUI) has been created in Tkinter ver. 8.5.9 for Python.

**Figure 1.**
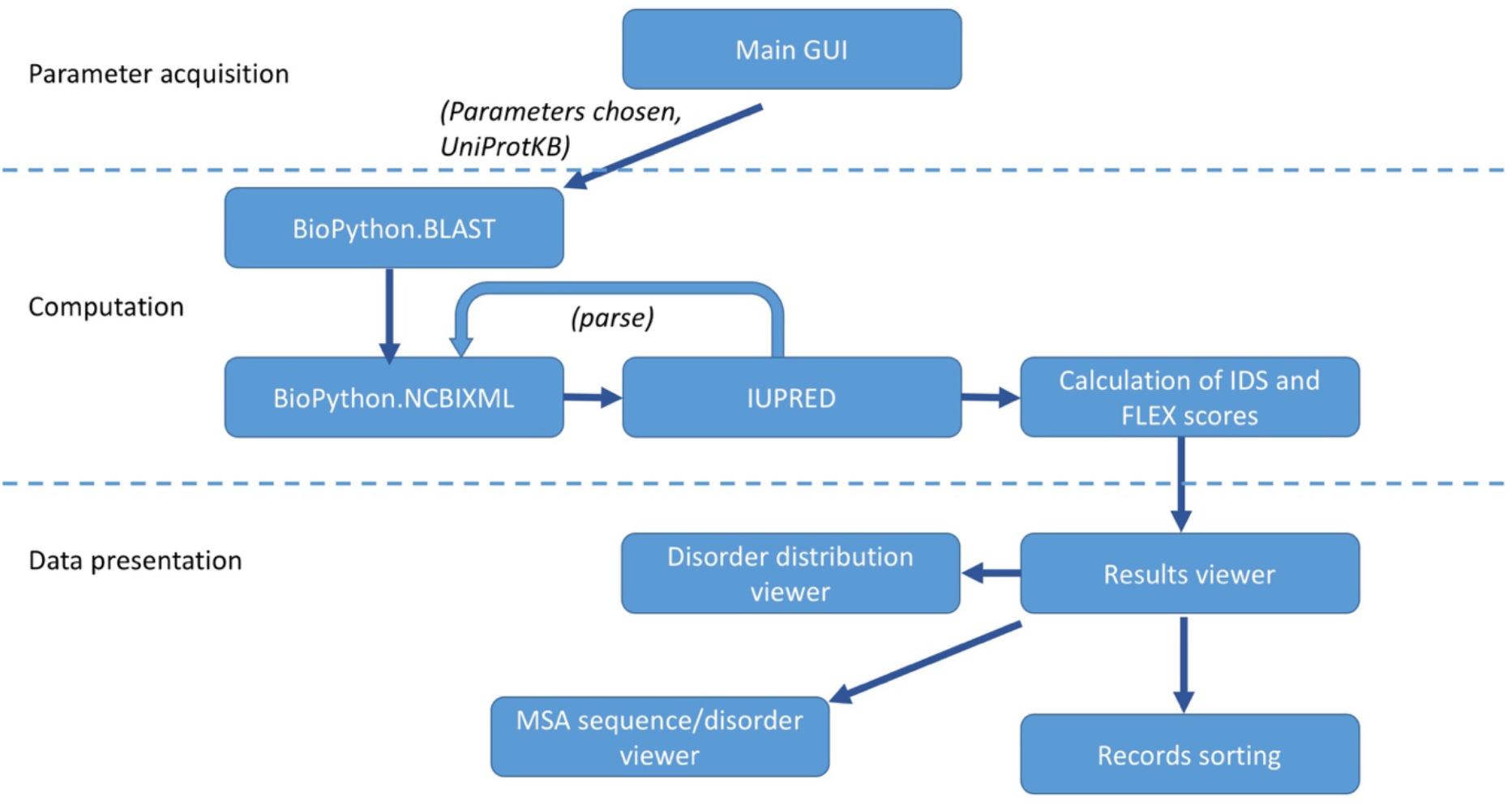
The component architecture of BASILIScan, classified into the three stages of: parameter acquisition, computation and data presentation.

### Sequence-based similarity search

After the plain unformatted amino acid sequence is provided, a BLASTP (18) search is conducted against the selected sequence database (UniprotKB/Swissprot being the default and recommended one (19)) and the results displayed according to the filters set by the user – such as an expect value (E-value) threshold or a maximum number of hits. The Bio.Blast functionality of Biopython is used to run on-line BLAST. The .xml output file is parsed with the NCBIXML function in Biopython.

Users may perform searches on custom, remote, FASTA-formatted libraries instead, by selecting the library file through the “Advanced properties/Select database” option. Remote search on the selected library is then performed by BLAST+ ver 2.7.1, including conversion of the library by makeblastdb. Sequence identifiers used in remote libraries should be Uniprot-derived in order for all BASILIScan modules to work correctly.

### Handling of viral polyproteins

In the case of the resulting sequences constituting a viral polyprotein, BASILIScan will perform virtual processing of the polyprotein and will apply all its analysis and metrics tools to the appropriate fragment only. This functionality is only offered when UniprotKB/Swissprot database is selected for search, due to lack of proteolytic processing information in non-manually curated databases. This option can also be disabled in “Edit/Advanced preferences”.

### Prediction of intrinsic disorder

For BLAST hits fulfilling the criteria set by the user, intrinsic disorder is calculated by the IUPRED algorithm (11), which has been used extensively in other publications for predicting intrinsic disorder *in silico* (20-23). Two alternative IUPRED modes are available: “long disorder” or “short disorder”. For most applications, the “long disorder” mode is suggested. The intrinsic disorder score (IDS) of each entry is then calculated in the following way:

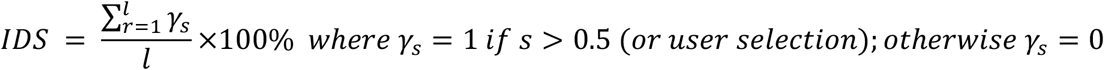

where *l* is the length of the protein and *s* is the residue’s IUPRED score.

### FLEX score computation

Since identification of a homologue with superior intrinsic disorder properties requires scoring at least two parameters simultaneously, the hybrid FLEX score has been implemented. The FLEX score incorporates a weighted average of the intrinsic order and a hyperbolic transform of the E-value parameter in the following fashion:

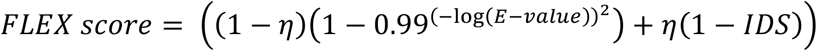

The weight is determined by the FLEX coefficient (*η*), which is set by the user before the homology search is run. Allowed values are between 0 and 1 and will shift the contribution ratio of intrinsic structural order (*1 - IDS*) to E-value transform. The hyperbolic transform of the E-value is meant to converge the extreme low-end E-value range while resolving the high-end and middle ranges. Consequently, the function of the logarithm of the E-value is sigmoidal, and bound from 0 to 1. It is characterised by a near-linear relationship between arguments corresponding to E-values of 10^-3^ and 10^-14^, while either tail approaches 0 or 1, respectively (supplementary figure S1).

### Visualisation of results

Results of a sequence query are presented in a table, for each hit showing the UniProt identifier, the GeneID, the expect value (E), sequence identity, similarity, the IDS score and the FLEX score. By default, the result hits are sorted from the lowest to the highest E-value. Sorting priority can be adjusted at any time from the main menu. The right-hand-side menu allows for more in-depth analysis of results. The ‘View’ option acquires the most important parameters of the selected item from the UniProt repository, displaying information such as sequence length, molecular mass and organism taxonomy.

The “Details” button draws a detailed trace of intrinsic disorder of the selected record within an interactive two-coordinate environment, implemented with Matplotlib. The environment allows for enlargement of selected parts of the trace, as well as for its translation. The option of exporting the graph as an image file is also provided. Traces can be overlayed on top of each other and therefore the intrinsic disorder can be explicitly compared between multiple protein records simultaneously.

Importantly, if the “enable disorder trace alignment” setting is switched on, the multiple disorder traces for the selected protein records will be automatically aligned on the axis, according to a multiple sequence alignment conducted in CLUSTALW (24). Default CLUSTALW parameters can be adjusted in Edit/Advanced properties. Any gaps inserted through the alignment algorithm will be visible in the aligned traces as residues with the IUPRED disorder score of 0.0 – an extremely unlikely occurrence for a protein residue otherwise.

### Distribution

Windows and OSX binary distributions of BASILIScan were packaged with Py2exe and Py2app, respectively. Packages for both platforms are freely available under the GNU distribution license and can be downloaded at www.basilisc.com/downloads. Open-source version is also available. Please consult the ReadMe file for further instructions on installation and running of BASILIScan, as well as for the dependencies required to run the open-source version (www.basilisc.com/readme/).

## Results and discussion

### Case scenario: human CDC7 protein kinase

Every BASILIScan job starts with providing a raw amino acid sequence and a job title. Next, the IUPRED mode is selected. Unless one is looking for very short stretches of disorder, or within very short protein sequences, the “long disorder” mode should be selected. The “E-value threshold” indicates the upper limit of the BLAST E-value, above which any hits will be ignored. The number of homology hits found can also be capped. Lastly, the “FLEX score priority coefficient” has to be set. This parameter will not influence which results are shown, but is meant to adjust the sensitivity of the FLEX score by shifting the contribution of homology versus structural order towards the final score. This means, for instance, that when the “priority coefficient” is set to 100% (all the way towards structural order), the FLEX score will only reflect the intrinsic order *(1-IDS)* of a given hit and will ignore the homology score component. The adjustment of the “priority coefficient” is particularly useful in cases where the BASILIScan run yields many hits containing one or more subpopulations clustered around particular values of IDS or E-value. The default priority coefficient value is 50%, in which case both terms will be taken into account equally.

To showcase the functioning of BASILIScan, the human cell division cycle protein kinase 7 (hCDC7, Uniprot identifier: 000311) was chosen as the sequence of interest. The protein is a 63.9 kDa serine/threonine protein kinase, which is an essential S-phase kinase implicated in DNA replication control and cell proliferation (25). The choice of hCDC7 kinase was dictated by availability of subject literature and structural information, as well as its functional conservation across many species. The 574-amino acid sequence of hCDC7 was submitted to BASILIScan for search in “long disorder” IUPRED mode, with an E-value threshold of 10^-10^, maximum number of hits of 100 and a FLEX score priority coefficient of 50%. Figure 2 and supplementary Table S1 show, respectively, the way results are displayed by the BASILIScan GUI and a full results dataset from the above search.

**Figure 2.**
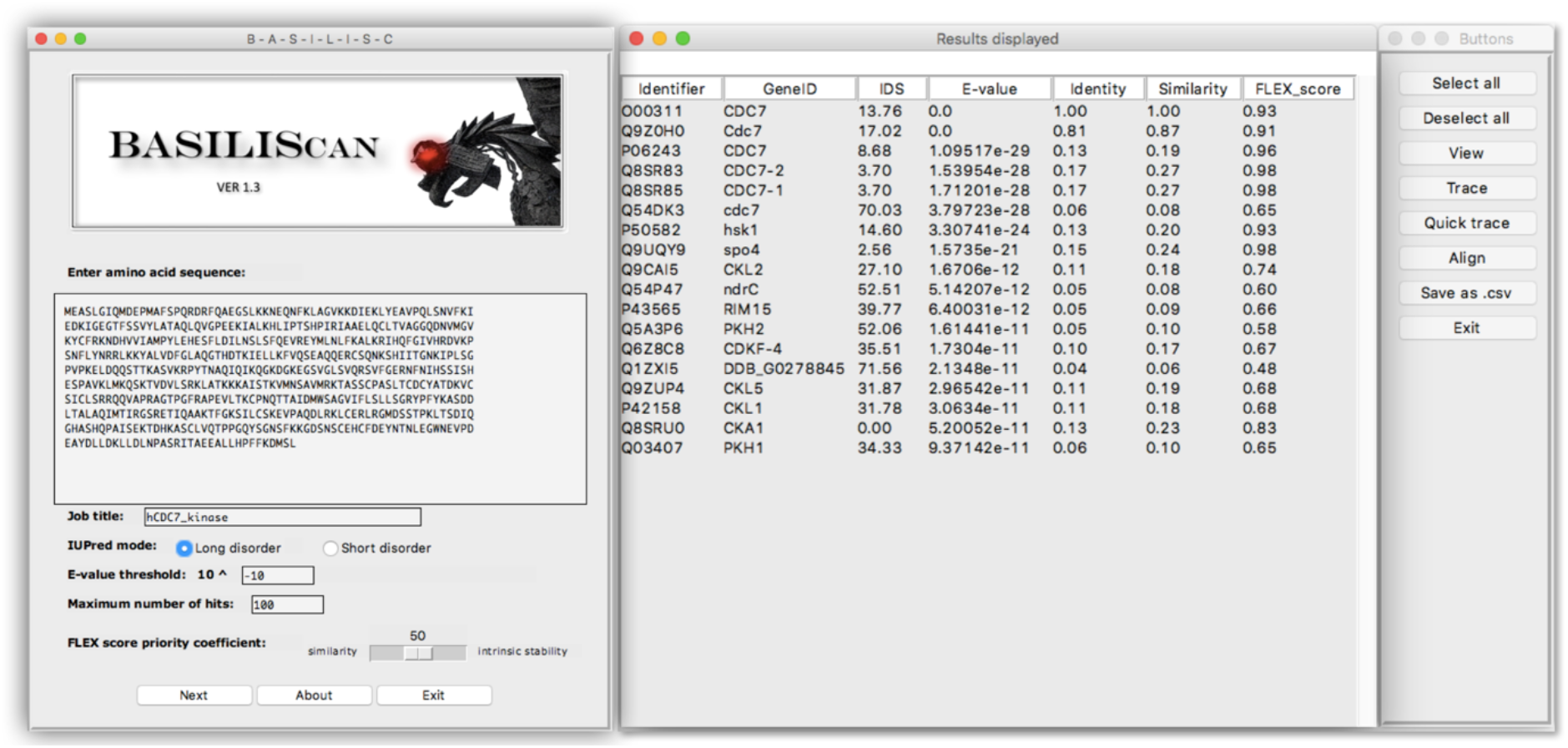
Presentation of search results by the BASILIScan GUI. The left panel provides a choice of basic parameters. Other parameters can be accessed through the “Advanced Properties” menu button. The middle panel shows the most important characteristics and calculated parameters of each result hit. Upon selection of one or more hit identifiers, the right panel buttons will activate different analysis modules.

The BASILIScan search yielded 18 hits, including the search query entry which always appears as the first row in the results window and always returns the E-value of 0. It is immediately apparent that although many homologues with low E-values were found, their intrinsic disorder varies considerably (from 0% to 72%). Human CDC7 kinase shows an IDS score of 14%, with many residues oscillating around the 0.5 disorder threshold. This can be seen by using the module “Trace” which graphs the disorder scores for every residue in a given sequence (Figure 3A). Indeed, lowering the IUPRED score calculation threshold from the default 0.5 to 0.4, leads to a dramatic increase in IDS of hCDC7 up to 35%. The FLEX score, being a weighted average of the E-value transform and the IDS score, should in most cases serve as the easiest measure of identifying suitable hits. The most promising hit returned by this BASILIScan search, possessing the highest FLEX score, is a probable CDC7 kinase homologue from a fungus *Encephalitozoon cuniculi.* In addition, a remote search with BASILIScan against a TrEMBL database encompassing all available vertebrate proteomes has also found numerous homologous sequences with IDS and FLEX scores superior to the search query statistics.

**Figure 3.**
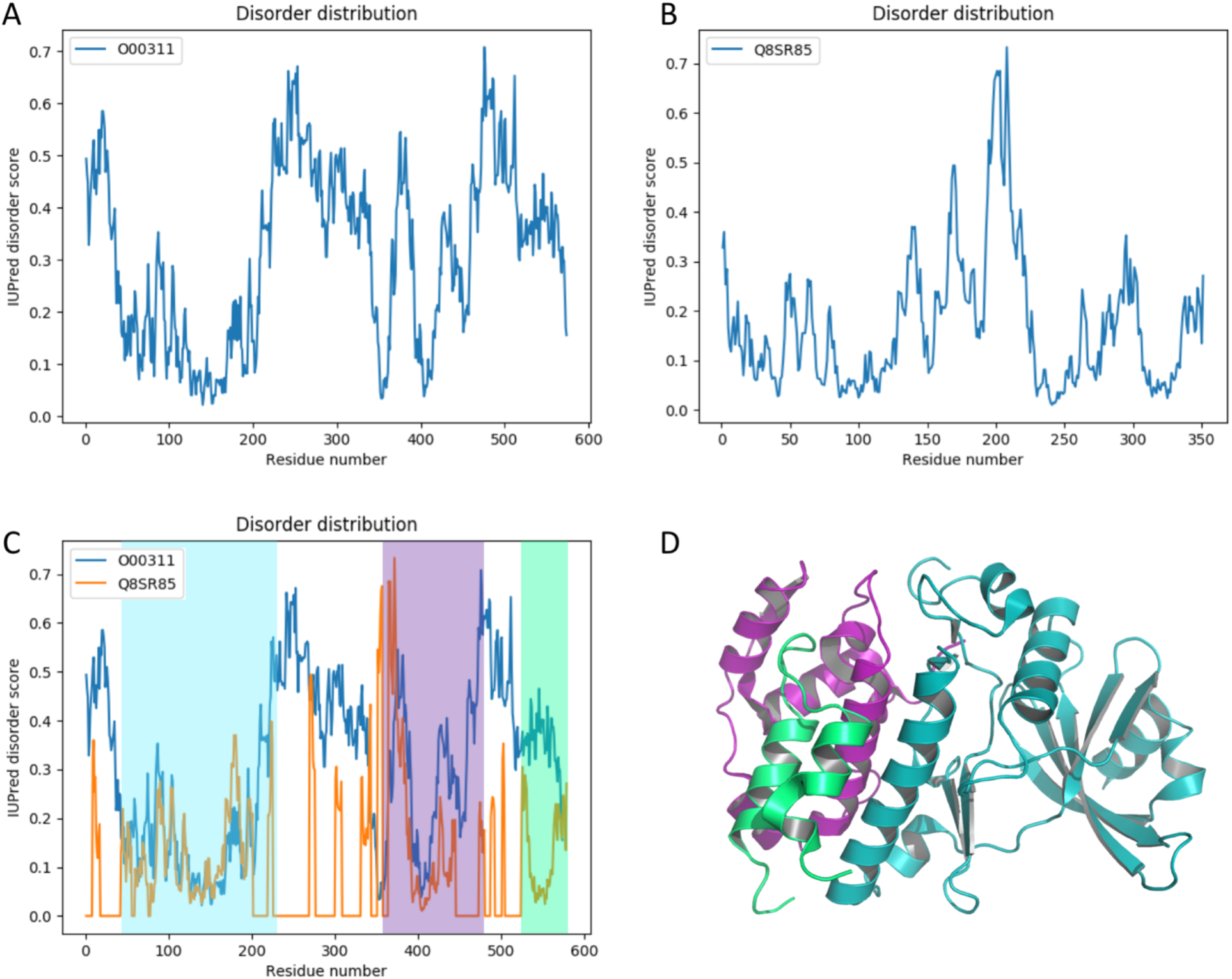
Detailed traces of putative intrinsic disorder drawn by the “Trace” module (A-C). Extensive regions of disorder are apparent from the disorder trace of human CDC7 kinase(A). The disorder trace of the highest-scoring hit in the BASILIScan search, the CDC7 kinase homologue of *E. cuniculi,* reveals a much more ordered appearance (B). Disorder trace alignment module allows for identification of the disordered regions in human CDC7 (blue trace) that are missing in the fungal homologue (orange trace). The resulting structured domains are highlighted in cyan, purple and green (C). A deletion construct of human CDC7 kinase composed of the regions highlighted in panel C has been successfully crystallised and its structure solved to 2.3 Å resolution (PDB: 4F99) (26) (D) Regions of the structure in panel D are coloured corresponding to the highlighting colour in panel C.

Overlaying the calculated intrinsic disorder trace of the submitted sequence (000311) on top of the BASILIScan hit of the highest FLEX score (Q8SR85), clearly shows the conservation of some intrinsic disorder patterns (residues 363-390) and confirms that there are regions of diminished intrinsic disorder (e.g. residues 420-446, 532-574) as well as significant deletions of particularly disordered segments (residues 1-40, 202-331) in the fungal and vertebrate homologues (Figure 3A-C). The “align” feature can also be used to perform a multiple sequence alignment on multiple selected hits and overlay the calculated intrinsic disorder scores for each residue in the alignment as a heatmap, in order to compare the conservation of disorder between many sequences at once (Figure S2).

The crystal structure of human CDC7 kinase has been reported (26). Crystallisation was achieved through identification of disordered regions by limited proteolysis and NMR spectroscopy, with their subsequent deletion. The crystallised protein encompasses the three ordered regions predicted by BASILIScan (highlighted in cyan, purple and green in Figure 3C-D).

### Occurrence of structured homologues of human proteins in other species

In order to learn about the scope of BASILIScan applicability in finding structured homologues of intrinsically-disordered proteins, every sequence of the human proteome (all 20243 sequences available in UniprotKB/Swissprot) was subjected to an individual BASILIScan search with the same parameters as used for hCDC7 above. All human sequences were removed from the results database. This high-throughput search resulted in disorder prediction and scoring of nearly 100,000 protein sequences, and showed that a surprisingly large extent of the human proteome has more intrinsically-ordered relatives in other species. BASILIScan identified at least one homologue exhibiting FLEX score higher than that of the query sequence for the vast majority of the human proteome (14865 sequences or 78%) and ten or more such homologues for over 30% (Figure 4A).

**Figure 4.**
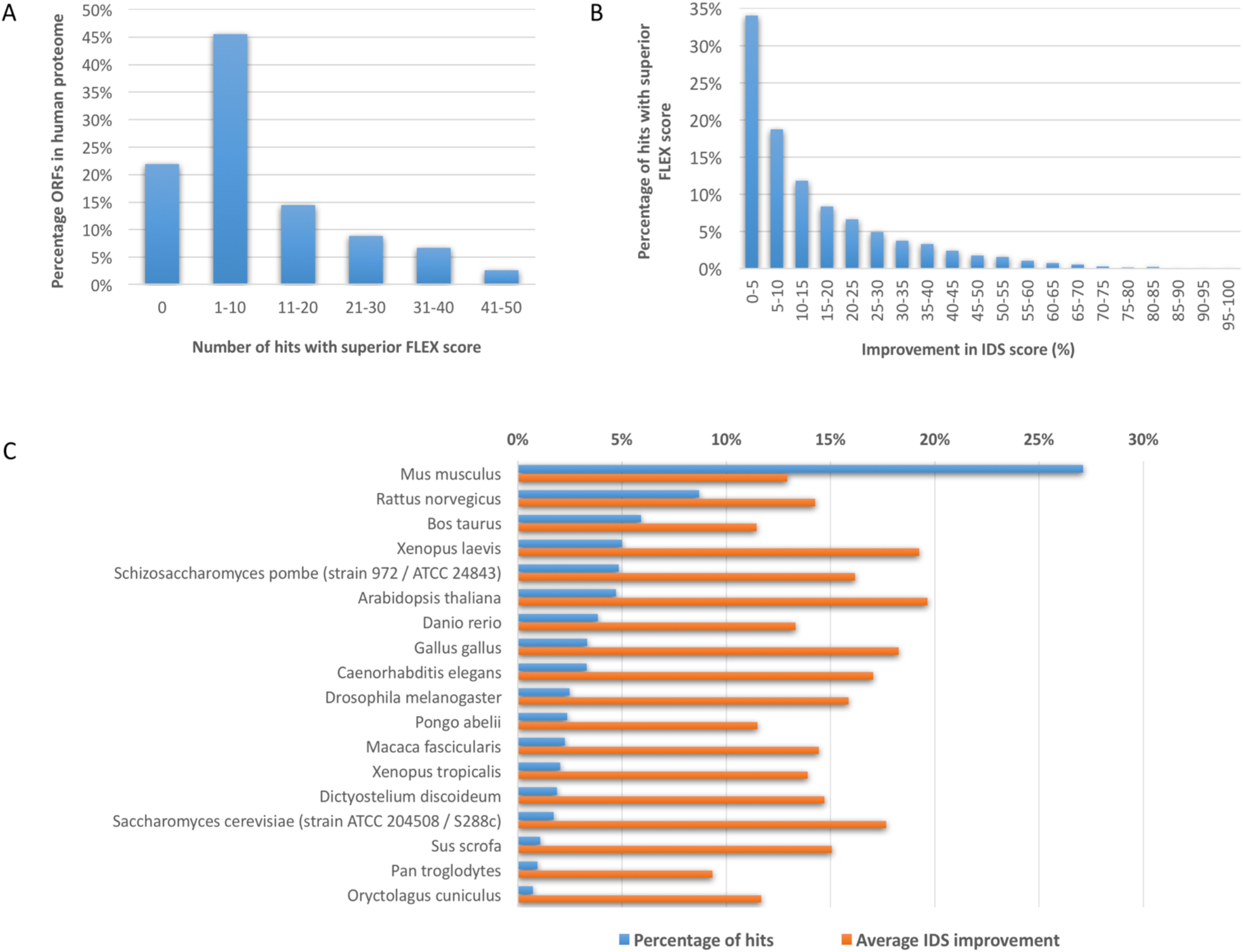
Results of a high-throughput BASILIScan search with every ORF in the human proteome. The success of this search can be assessed by the number of homologous sequences found with a FLEX score higher than that of the query sequence (A). Limiting the results dataset to the queries which yielded at least one such homologue, the extent of improvement in IDS score can be plotted in 5% increments (B). Information about the most common species of the highest-scoring homologue for each query was also plotted (blue), along with the corresponding average difference in IDS (orange).

Importantly, in just under half of hits scored as promising – possessing FLEX scores higher than the query – the improvement of intrinsic disorder represented by the difference in IDS score was very significant. The IDS score improvement of 10% or more was found in 47% of such BASILIScan-identified hits, while 27% of hits showed at least 20% improvement (Figure 4B). The top hits resulting from the BASILIScan search belonged to 631 unique species, but by far the highest occurrence was displayed by mouse homologues (Figure 4C). The improvement in IDS score appeared not to correlate with the hit frequency of occurrence in the species of origin.

## Conclusions

As structural biology is evolving from tackling thermodynamically-stable, well-folded protein domains into much larger, multimeric and multicomponent protein systems, the need for rapid assessment and minimisation of intrinsic disorder becomes increasingly important. Software developed to assist recombinant expression construct design for maximising protein stability and crystallisibility through computing of parameters sourced from previously-solved crystal structures (XtalPred, (27)) and surface entropy reduction (SERp, (28)) have been used extensively. However, to the author’s knowledge, user-friendly and widely-accessible tools for high-throughput comparative analysis of intrinsic disorder patterns in related proteins have not been developed and explored for the purpose of recombinant expression construct design.

The functionalities implemented in BASILIScan serve two main experimental purposes: identification of homologues with diminished intrinsic disorder; and recognition of disordered regions suitable for deletion due to limited conservation. For both applications, the accessible user interface and the adjustable parametrisation are invaluable for identification of the most promising candidates and regions. As demonstrated by the high-throughput BASILIScan screen of the human proteome, identification of a more intrinsically-stable homologue of an IDP is likely to be feasible in most cases. Therefore, BASILIScan should be a valuable and easily-accessible resource for more streamlined rational expression construct design approaches and study of IDPs.

## Availability of data and material

Ready-to-use binary distributions of BASILIScan (currently version 1.3) are available for free for academic users. Packages for OSX and Windows systems can be downloaded from the project website: www.basiliscan.com/download. Open-source Python code will be released through the website upon publication. Instructions on which dependencies need to be installed for compiling the open-source version are under www.basilisc.com/readme. Please also consult the website for tutorials, FAQs and updates.

All database files generated in this publication will be available for download from www.basiliscan.com upon publication of the paper.

## Acknowledgments

I would like to acknowledge Dr Goedele Maertens as well as Mr Michal Kosicki, Dr George Gerogiokas and Dr Robert White for fruitful discussions, suggestions, a critical review of the manuscript and testing of the initial versions of BASILIScan.

## Competing interests

The author declares no competing interests.

## Supplementary information

**Table S1.** Summary of BASILIScan results for homologues of human CDC7 kinase (Uniprot/Swissprot identifier 000311). Search parameters and the scoring system are described in “Results”. Calculations of the IDS and FLEX score parameters are described in the “Design and Implementation” section.

| *Identifier* | *GeneID* | *IDS* | *E-value* | *Identity* | *Similarity* | *FLEX_score* |
| --- | --- | --- | --- | --- | --- | --- |
| **O00311** | CDC7 | 13.763 | 0 | 1.000 | 1.000 | 0.931 |
| **Q9Z0H0** | Cdc7 | 17.021 | 0 | 0.809 | 0.874 | 0.915 |
| **P06243** | CDC7 | 8.679 | 1.10E-29 | 0.134 | 0.191 | 0.956 |
| **Q8SR83** | CDC7-2 | 3.704 | 1.54E-28 | 0.174 | 0.268 | 0.981 |
| **Q8SR85** | CDC7-1 | 3.704 | 1.71E-28 | 0.174 | 0.268 | 0.981 |
| **Q54DK3** | cdc7 | 70.028 | 3.80E-28 | 0.057 | 0.083 | 0.650 |
| **P50582** | hsk1 | 14.596 | 3.31E-24 | 0.128 | 0.195 | 0.925 |
| **Q9UQY9** | spo4 | 2.564 | 1.57E-21 | 0.147 | 0.235 | 0.981 |
| **Q9CAI5** | CKL2 | 27.097 | 1.67E-12 | 0.110 | 0.176 | 0.740 |
| **Q54P47** | ndrC | 52.509 | 5.14E-12 | 0.050 | 0.079 | 0.599 |
| **P43565** | RIM15 | 39.774 | 6.40E-12 | 0.050 | 0.089 | 0.659 |
| **Q5A3P6** | PKH2 | 52.059 | 1.61E-11 | 0.052 | 0.102 | 0.585 |
| **Q6Z8C8** | CDKF-4 | 35.512 | 1.73E-11 | 0.105 | 0.174 | 0.666 |
| **Q1ZXI5** | DDB_G0278845 | 71.556 | 2.13E-11 | 0.037 | 0.057 | 0.483 |
| **Q9ZUP4** | CKL5 | 31.871 | 2.97E-11 | 0.113 | 0.189 | 0.677 |
| **P42158** | CKL1 | 31.778 | 3.06E-11 | 0.109 | 0.178 | 0.676 |
| **Q8SRU0** | CKA1 | 0.000 | 5.20E-11 | 0.125 | 0.226 | 0.827 |
| **Q03407** | PKH1 | 34.334 | 9.37E-11 | 0.059 | 0.097 | 0.646 |

## Supplementary figures

**Supplementary figure S1.**
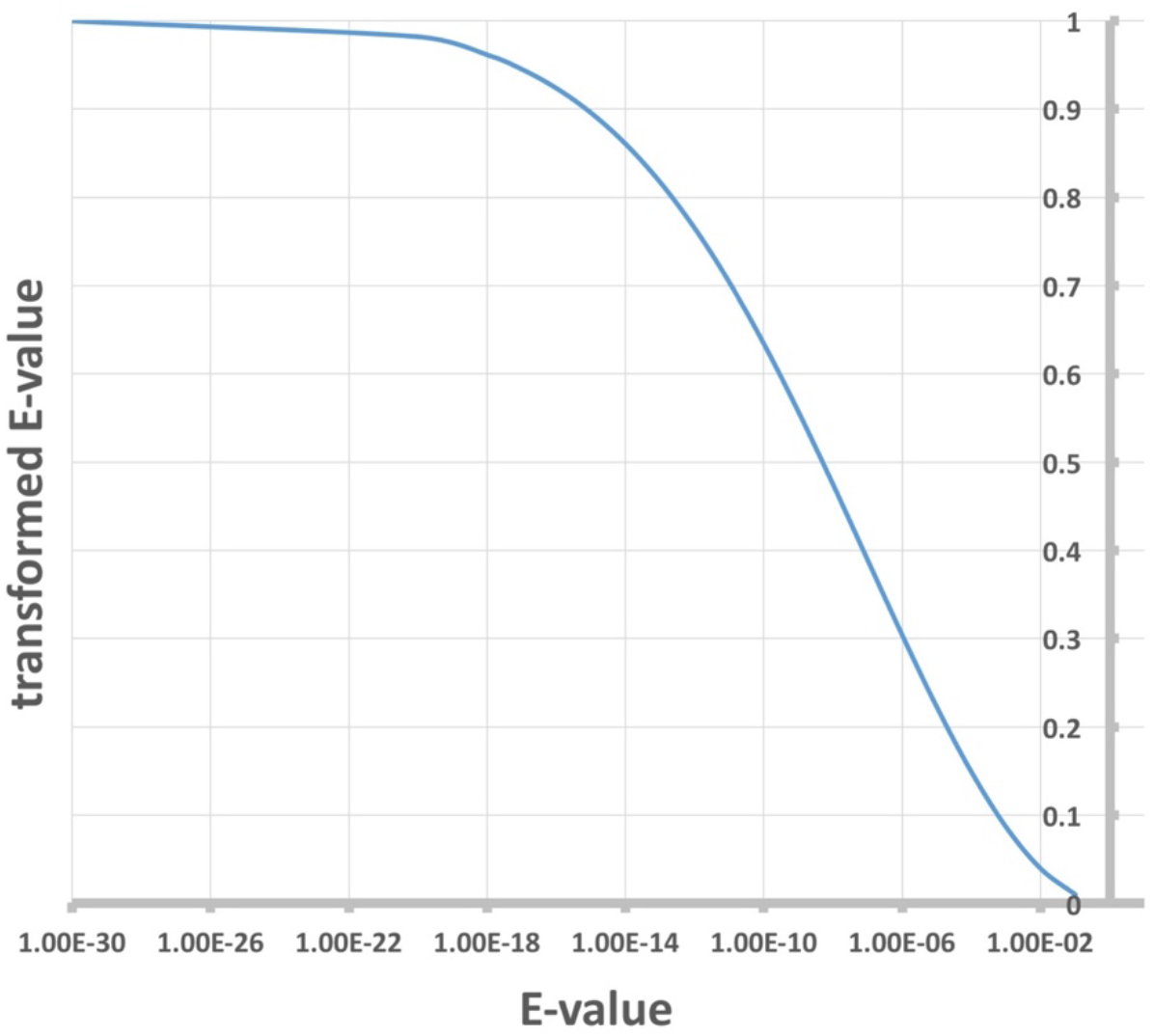
For more accurate scoring purposes, a hyperbolic transform is applied to the BLAST-derived E parameter, resulting in a value bound from 0 to 1 for E-values of below 1. The function is near-linear for E-values of between 10^-14^ and 10^-3^, while both far ends reach a plateau. The transformed E-value is then used for calculation of the FLEX score.

**Supplementary figure S2.**
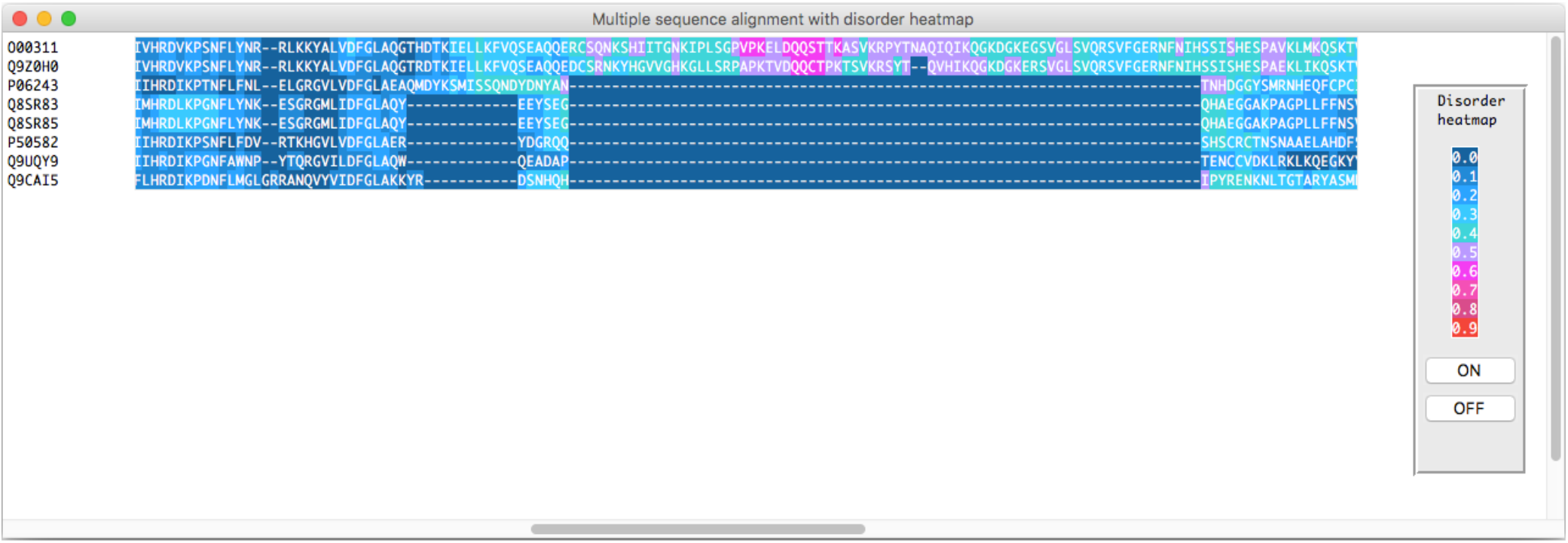
Eight highest-scoring hits from a BASILIScan search against hCDC7 kinase (Uniprot/Swissprot identifier 000311) aligned with the “Align” module and with the calculated intrinsic disorder overlayed on the alignment as a heatmap. In this case, the absence of an extended, highly-disordered fragment in the bottom five sequences is apparent.

